# “Viscotaxis”- Directed Migration of Mesenchymal Stem Cells in Response to Loss Modulus Gradient

**DOI:** 10.1101/804492

**Authors:** Pallavi Uday Shirke, Hiya Goswami, Vardhman Kumar, Darshan Shah, Siddhartha Das, Jayesh Bellare, K.V. Venkatesh, Jyoti R. Seth, Abhijit Majumder

## Abstract

Directed cell migration in response to chemical and mechanical gradients plays a crucial role in physiological and pathological conditions. One such mechanical cues that is known to influences cell migration is the gradient of substrate elastic modulus (*E*). However, the elastic modulus alone cannot fully define the material properties of the cellular microenvironment, which often has both elastic and viscous characteristics. In this study, we investigated the influence of the gradient of viscous nature, as defined by loss modulus, G”, on cell migration. We cultured human mesenchymal stem cells (hMSCs) on a collagen-coated polyacrylamide gel with constant elastic property, as defined by the storage modulus G’, but with the gradient of loss modulus G”. We found hMSCs to migrate from high to low loss modulus. We have termed this, thus far unreported, directional cellular migration as “Viscotaxis”. We have confirmed uniform collagen density and constant storage modulus of the gel by fluorescence microscopy and atomic force microscopy to eliminate the possibilities of haptotaxis and durotaxis. We hypothesize that material creep in the high loss modulus region hinders the building up of the cellular traction, leading to a force asymmetry that drives the observed viscotaxis. To verify our hypothesis, we estimated the cellular traction on gels with high and low loss moduli. We indeed found that cells apply higher traction force on more elastic materials i.e. materials with low loss modulus. On the disruption of actomyosin contractility with myosin inhibitor blebbistatin and ROCK inhibitor Y27632, directional migration was lost. Further, we showed that cells can maintain a stable morphology on a low loss modulus substrate but due to its inability to build up stable cellular traction on a substrate with high loss modulus, the cell spreading remains in a dynamic state. Our findings in this paper highlight the importance of considering the viscous modulus while preparing stiffness-based substrates for the field of tissue engineering.

## Introduction

The directed cell migration or “taxis” in response to different microenvironment cues, is crucial during development^1^, wound repair^2^, and immune responses^3^. Depending on the cue that is responsible for directed migration, several different types of “taxis” have been reported in the literature such as chemotaxis^4^, haptotaxis^5^, rheotaxis^6^, curvotaxis^7^, topotaxis^8^, mechanotaxis^9^ etc. Many of such reported “taxis” are based on mechanical cues^10, 11^. In 2000, Lo *et. al*. demonstrated that cells migrate from high to low Young’s modulus region on a substrate with a rigidity gradient. This process is known as “durotaxis”^12^. However, Young’s modulus alone cannot completely describe the biological materials such as cells, matrices and tissues, which are known to be viscoelastic^13–15^. Yet surprisingly, there are only a handful of literature that have studied the effect of both storage modulus G’ and loss modulus G”, on cell functions. Those studies demonstrated that viscoelasticity of the substrate influences cellular phenotype, differentiation, and migration ^16–18^. For example, substrates with high loss modulus influence mesenchymal stem cells (MSCs) to differentiate into the myogenic lineage whereas elastic substrate directs them towards osteogenic differentiation^19^. Also, stress relaxation of the substrate was shown to enhance cell spreading^20, 21^. Substrate viscoelasticity has been shown to influence cellular migration as well. On high loss modulus substrate, the velocity of MSCs is significantly higher than on low loss modulus substrate^22^. Substrate viscoelasticity also enhances the correlation between the movement of cells in an epithelial sheet^23^. A recent study has shown that fibroblast cells show amoeboid migration on high loss modulus substrate due to weak substrate adhesion and low stress fiber formation. However, on more elastic substrates they adhere strongly with more stress fibers leading to the mesenchymal mode of migration^24^.

While the reported literature discusses the effect of uniform viscoelasticity on cell migration, the *in vivo* microenvironment is rarely uniform. To address how the cells respond to a gradient of loss modulus, in this paper we created substrates with a gradient of loss modulus and observed the migration of human mesenchymal stem cells (hMSCs) on them. First, we extensively explored the rheological properties of polyacrylamide gels (PAA gels) made from different combinations of acrylamide and bisacrylamide. Based on the results, we selected two gel compositions with the same storage modulus but different loss moduli. Using these two combinations we created gels with a loss modulus gradient but storage modulus constant. We found that cells preferentially migrate from high loss modulus to low modulus region. Further, an increase or decrease in the gradient strength resulted in an increase or decrease in migration bias respectively. Inhibition of actomyosin contractility by pharmacological inhibitors disrupted this preferential migration making cells nonresponsive to the loss modulus gradient. Also, higher membrane fluctuation of HeLa cells on high loss modulus substrate suggest dynamic nature of cells compared to stable cell on low loss modulus. In summary, our study demonstrates that cells respond to loss modulus gradient and migrate from high loss to low loss modulus region causing directed cell migration which we are addressing as “*viscotaxis.*”

## Results

### Rheological characterization of polyacrylamide gel to study the effect of loss modulus gradient on cell migration

To study the effect of the gradient of loss modulus on cell migration, the primary challenge was to find acrylamide and bis-acrylamide concentrations that would give rise to polyacrylamide gels with similar storage modulus but different loss moduli. While many published literatures estimate Young’s modulus of PAA gel with predefined acrylamide and bis-acrylamide ratio, only limited data are available in the public domain describing their complete rheological properties ^19, 20^. To bridge this gap, we prepared PAA gels with varying acrylamide and bis-acrylamide concentrations and characterized the rheological properties of the resulting PAA gels, as shown in table S1. From this table we selected two compositions that give rise to gels with the same storage modulus but widely different loss moduli (Table 1). Time sweep oscillatory rheometry was performed to determine the gelation time of the polyacrylamide gel (Fig.S1). After confirming the crosslinking of these two PAA gels, we performed frequency sweep 100 to 0.01 rad/s (Fig.S2). In this paper, we will call the gel with high loss modulus as *High Loss* (G’ =1.2 ± 0.12 kPa, G”= 289 ± 18 Pa) and the gel with low loss modulus as *Low Loss* (G’ =1.2 ± 0.05 kPa, G” = 30 ± 6.13 Pa) (Fig 1a). The pre-polymer compositions that give rise to *High Loss and Low Loss* gels will be termed as High Loss Composition (HLC) and Low Loss Composition (LLC).

**Fig. 1.**
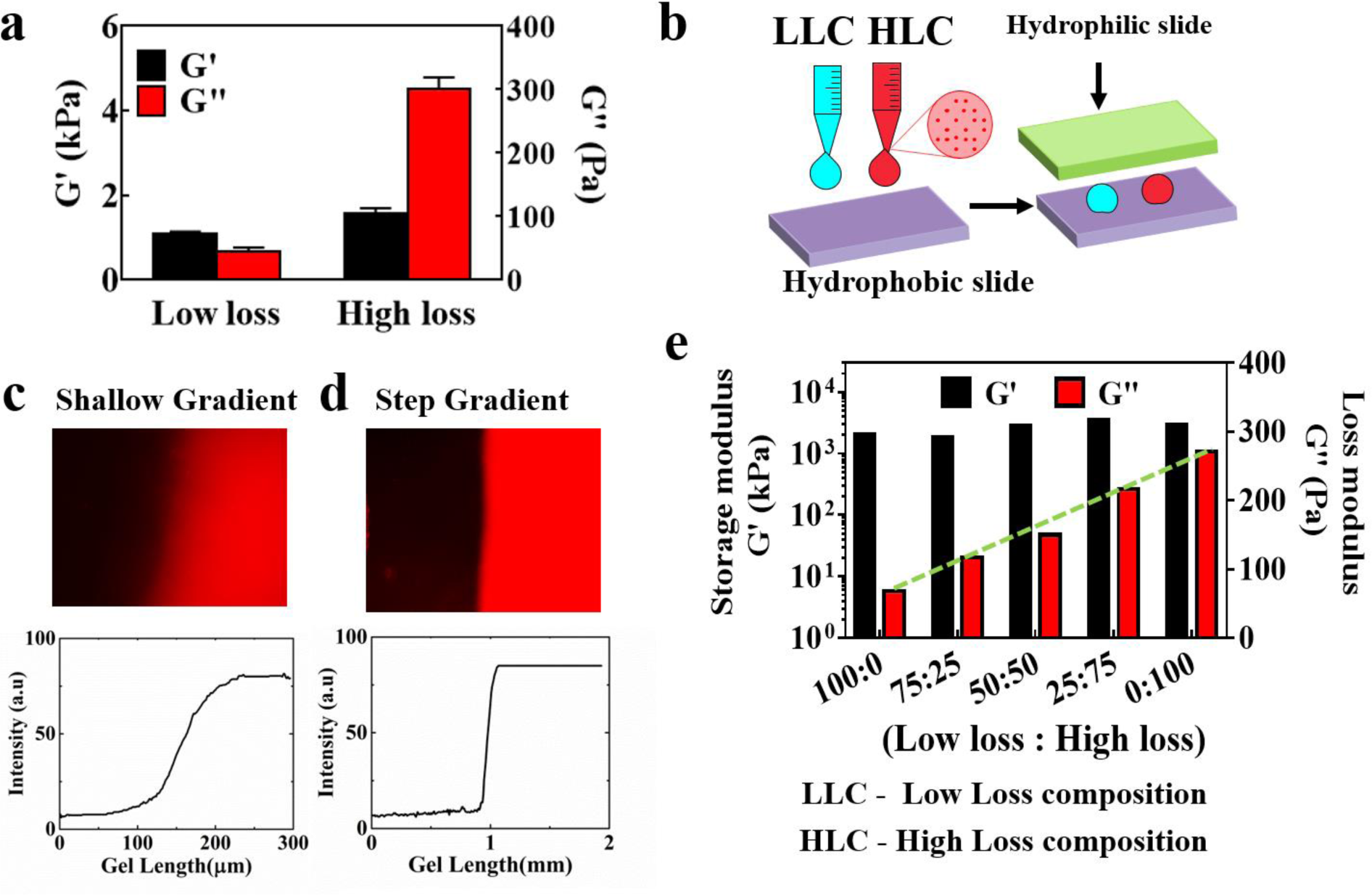
Generation of substrates with loss modulus gradient and their characterization. (a) Storage Modulus (G’) and Loss Modulus (G”) at 0.01 Hz frequency of Low Loss and High Loss substrates. (b) Schematic of the loss modulus gradient generation. Fluorescent red polystyrene bead intensity measured along the (c) shallow loss modulus gradient and (d) step loss modulus gradient gels with respective fluorescent intensity plotted against the gel length. (e) The storage modulus (G’) and loss modulus (G”) of samples with LLC: HLC such as 100:0, 75:25, 50:50 and 25:75, 0:100.

**Table 1:** Polyacrylamide gel samples selected for the study along with their composition and mechanical properties.

| Sr. No. | Acryl % | Bis % | Storage Modulus<br>(kPa) | Loss Modulus<br>(Pa) | Sample Name |
| --- | --- | --- | --- | --- | --- |
| 1 | 6.68 | 0.06 | $1.2 \pm 0.12$ | $30 \pm 6.13$ | Low Loss |
| 2 | 8 | 0.26 | $1.5 \pm 0.05$ | $289 \pm 18$ | High Loss |

To prepare the gel with a gradient of loss modulus, we used two drop technique as reported earlier^12 25^. In short, we placed two drops of pre-polymer solutions with two different compositions, HLC and LLC, at proximity on a hydrophobic coverslip (Fig. 1b). Next, we placed a hydrophilic plate on top to make the drops coalesce and diffuse into each other. Two competing processes run here in parallel; diffusion and polymerization. Depending on the relative dominance of these two processes, the strength of the gradient is established. We added red fluorescent polystyrene beads of 0.2 µm into High Loss solution which helped us to visualize and quantify the gradient strength. Depending on whether we coalesce the drops and start the diffusion simultaneously with polymerization, or allow the polymerization to go for 5 min before coalescence, we get gels with two different gradient strengths. In the first process, the loss modulus changes gradually over a defined length. We call this gel as Shallow Grad substrate (Fig.1c). The second process gives a us a step change in the loss modulus. We name this gel as Step Grad substrate (Fig.1d). The change in this gradient is quantified by measuring the fluorescent intensity along the gel length.

Further, to ensure that the diffusion driven variation whilst pre-polymer composition along the gel length gives gradually changing loss modulus keeping the storage modulus unchanged, we prepared gels from solutions of different ratios of HLC and LLC. Rheometry data, as shown in Fig. 1e, confirms that the loss modulus indeed depends linearly on LLC: HLC ratio.

### hMSCs migrate in response to loss modulus gradient

To answer if the gradient of loss modulus induces directed cell migration, we recorded the movement of hMSCs on substrates with loss modulus gradient. We used uniform High Loss and Low Loss gels as controls. Movement of cells on these substrates was recorded using time-lapse microscopy for 18 h with 30 min time intervals. The videos were analyzed using “Chemotaxis Migration Tool” plug-in in ImageJ software. Fig 2a-c show representative rose plots. In Fig 2a, 18 cells (red lines) moved from High Loss to Low Loss region in the 18 h of observation, while only 4 cells (black lines) moved in the opposite direction. We term this, so far unreported, biased migration in response to the substrate Loss Modulus gradient as *Viscotaxis*. Such biased migration was absent on gels with uniform loss modulus, used as control (Fig 2b, c) (Video S1, S2 and S3).

**Fig.2.**
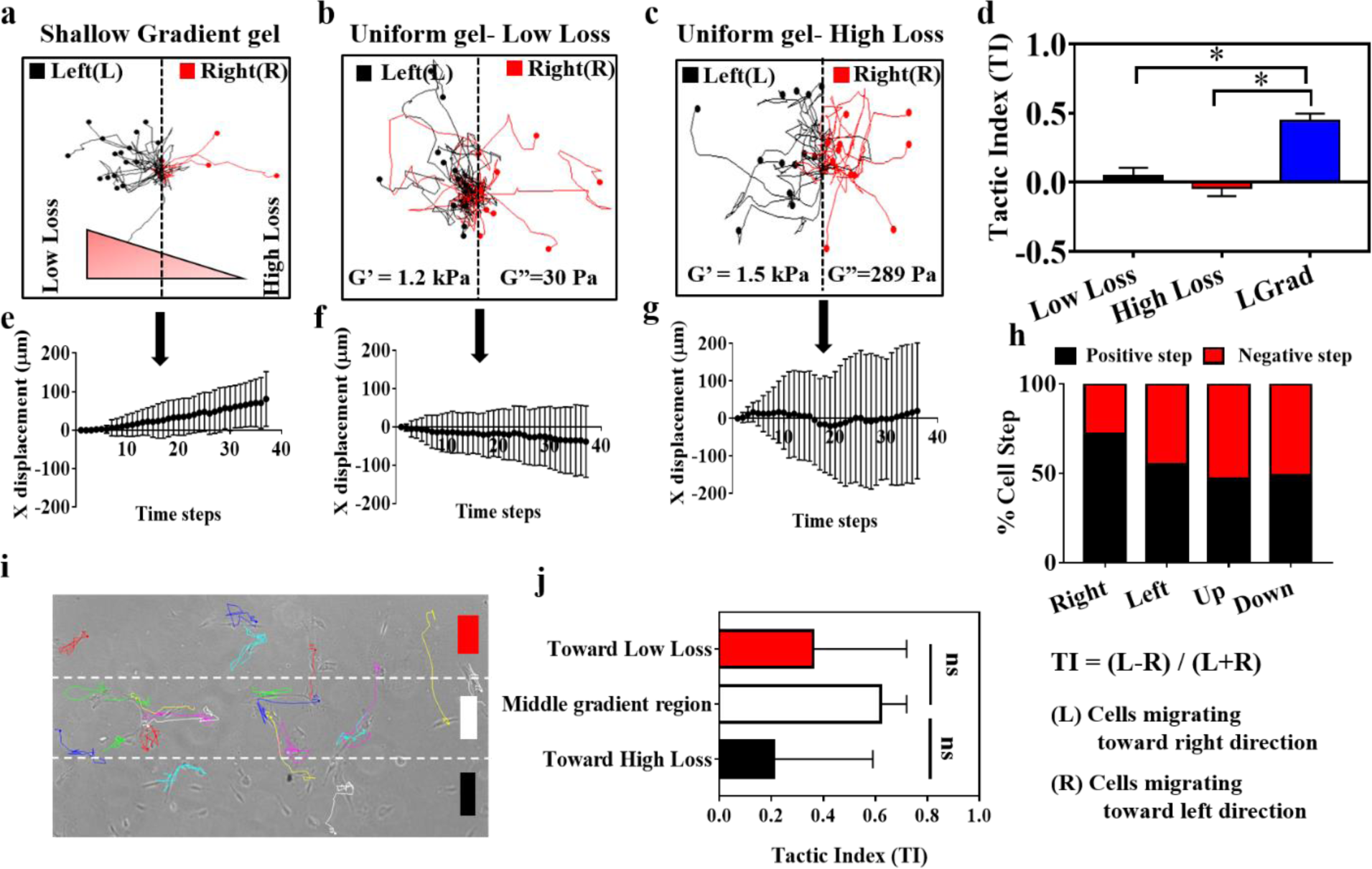
Viscotaxis, biased cell migration in response to a shallow gradient of loss modulus (G”). (a) Representative rose-plots depicting the trajectories of hMSCs migrating cells on the gradient of Low Loss – High Loss (shallow loss modulus gradient) (b) Low Loss modulus substrate (c) High Loss modulus substrate. (d) Tactic index at the end of 18 h for cells on shallow loss modulus gradient substrate, High Loss and Low Loss substrate. Evolution of tactic index from 0 h to 18 h with 30 min interval for cells on (e) loss modulus gradient gel (f) Uniform – Low Loss gel (g) Uniform- High Loss gel. (h) Quantitative analysis of cell steps (positive and negative step) migrating in four directions- right, left, up and down of the loss modulus gradient substrate. (i) hMSCs cells seeded on a loss modulus gradient. Dashed lines indicate three regions of the gradient region represented by red, white and black colour respectively. (j) Tactic index for the cells whose initial position falls in the three regions toward Low Loss, Middle and toward High Loss region. Error bar represents standard error mean (n = number of cells where * p< 0.05). Low loss substrate n = 107 cells, High Loss substrate n = 98 cells, shallow loss modulus gradient n = 232 cells for three independent experiments.

To further quantify the bias in cell migration, we estimated tactic index (TI), defined as the ratio of the difference in the number of cells moving in the two opposite directions to the total number of cells^25^. Mathematically,

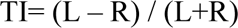

Where R and L are the numbers of cell migrated toward the right i.e., the high loss modulus region and to the left i.e. the low loss modulus region, respectively. Cell population with no bias, i.e., with equal probability of going toward either direction, has TI = 0. We found (Fig.2d) TI = 0.45 ± 0.05 on gradient substrate indicating that on an average 73% cells responded to the loss modulus gradient and migrated from High Loss to Low Loss region. It must be noted that, this fraction remains stable over time indicating that 30% of the cells are non-responsive. Cells on gels with uniform High Loss and Low Loss had TI = (0.04 ± 0.05) and (-0.03 ± 0.05) respectively, indicating no preference for directional migration.

While TI captures the overall bias in migration as a ratio, it fails to estimate the strength of the bias. In other words, TI calculation only considers the relative shift from the initial position for a cell but does not take into account the extent of shift. To overcome this limitation, we calculated the average x displacement over time. An average x displacement close to zero indicates random migration while the width of the distribution estimates the randomness.

Cells on the substrate with a loss modulus gradient showed a steady increase in x direction displacement. Also, it can be seen that bias migration took approximately 6 h to get established (Fig.2e).

Although in the case of uniform Low Loss, the average x displacement shows a slight deviation from zero, the standard deviation (SD) is distributed in both +x and –x region. In contrast, average x displacement on gradient gel, SD are all within the +ve x direction indicating steady bias. It must be noted that, SD on uniform High Loss modulus, is much larger compared to uniform Low Loss gel indicating more randomness in cell migration as compared to that on High Loss (Fig.2f, g).

From the TI data, we see that approximately 30% of the cells show negative viscotaxis, i.e., they migrate toward high loss region. We ask the question if this is an outcome of random walk or a decision of individual cell at every time point, we classified the steps into +ve steps and –ve steps. When a cell takes a step in the direction of decreasing loss modulus i.e., in the direction of Viscotaxis, we consider that step as +ve and when the step is in the opposite direction, we consider that step as –ve step. For analysis, we randomly chose 27 cells of 232 cells that showed +ve displacement after 18 h in our TI study. We call this population as viscotactic cells and abbreviate them as VisC. Similarly, we also randomly selected 27 cells which showed –ve displacement after 18 h in TI study. We call this population as non viscotactic cell or (NVisC).

We found that VisC took +ve steps in 77% of the cases (Fig. 2h). In contrast NVisC had no bias and took an almost equal number of +ve steps (55%) and -ve step (45%). To use as internal control, we analyzed the movement of the cells in the up down direction. For cells with net movement in up direction, +ve and -ve steps were 46% and 54% respectively. Cells with net downward migration took 49% +ve steps and 51% -ve steps. As expected, there was no bias in the up-down direction.

Next, we asked whether the initial location of a cell on a gradient determines its decision. In other words, we asked how the loss modulus gradient experienced by a cell at an initial time period influenced the migratory decision. To do that, we divided our field of observation into three parallel rectangular regions, each of width 130 μm (Fig.2i).

The regions close to the Low Loss side was named as zone 1. Similarly, the middle section was named as zone 2, and the region towards High Loss side was named as zone 3.

We found that in the zone 2, TI was 0.62 ± 0.25. In zone 1 and 3, TI= 0.45 ± 0.32 and 0.36 ± 0.62 respectively (Fig.2j). Though the mean TI was different for these three zones, the difference was not significant (p = 0.06). This indicates that the cell’s initial position within the frame of observation did not influence its migration bias. Together, these results highlight the possibility of a subpopulation of cells that are non-responsive to the given gradient strength.

### Viscotaxis depends on the loss modulus gradient strength

Next, to investigate the influence of the strength of loss modulus gradient on migration bias, we prepared two sets of gels. In the first case, we increased the loss modulus gradient strength by creating a step increase from G” = 30 Pa to G” = 300 Pa (Fig 3a-b). Whereas in the second case, we reduced the loss modulus gradient strength to 0.06 Pa/µm from 0.12 Pa/µm as used in the previous section. This was achieved by using two pre-polymer solutions, one that gives rise to a gel of G”=30 Pa and the other that gives rise to a gel of G”=150 Pa i.e., High (50:50 composition) Loss (Fig.1e).

**Fig. 3.**
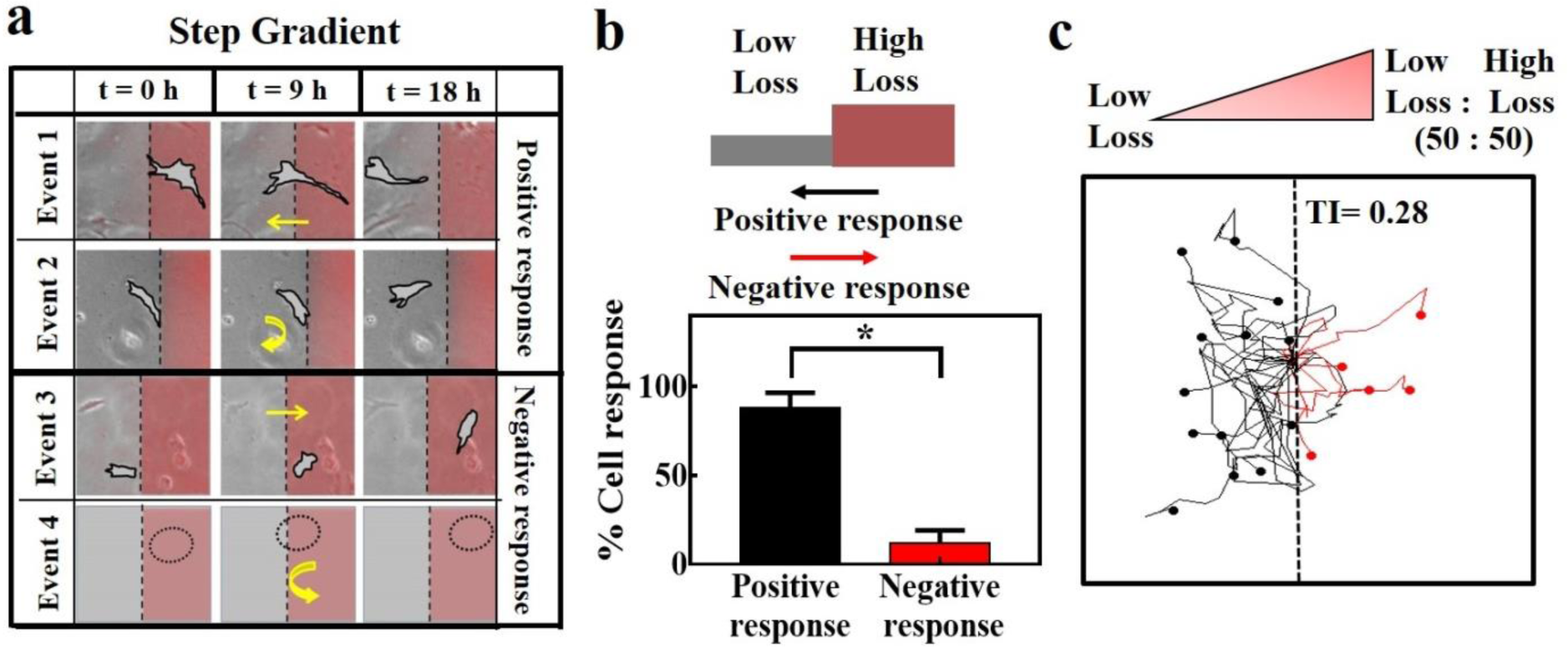
Effect of gradient strength on viscotaxis: (a) Classification of cells based on events at the interface of the two gel regions in case of Low Loss substrate – High Loss substrate gradient and a representative image for each event. (b) Representation of the percentage of cells responding to step gradient of loss modulus substrate. (c) Representative rose-plots depicting the trajectories of migrating cells on the gradient of Low Loss – High (50:50 composition) Loss.

In case of a step increase in loss modulus, the cells can take four probable trajectories, as shown in Fig.3a. Here, the top panel shows the possible event 1, in which the cell starts from the High Loss modulus region and crosses the boundary to reach the Low Loss modulus region (Video S4). In the possible event 2 (2^nd^ panel), the cell starts from the Low Loss modulus region, tries to cross the border but eventually takes a U-turn and returns to the Low Loss modulus region (Video S5). We group these two events as a positive response because the movement follows the expected Viscotaxis direction (preferred movement from High Loss to Low Loss modulus region).

The negative events, i.e. in which cellular movements contradict the predicted Viscotaxis direction are shown in panels 3 and 4. The third panel shows event 3 where a cell crosses over from Low Loss to High Loss modulus region (Video S6). The last panel shows the possible event 4 where a cell starts from High Loss modulus, approaches the Low Loss region but takes a U-turn from the boundary. In our time-lapse video done for four independent experiments recorded over 18 h, we could find 14 incidents of event 1, 23 incidents of event 2, and 9 incidents of event 3. Interestingly, we could not find any incident of event four. That is why, in Fig 3a, event 4 is presented with an imaginary cell with a dotted circle. Overall, we found that 85% of the cells show positive response demonstrating Viscotaxis (Fig 3b). In terms of TI, this percentage corresponds to as high as 0.7. In contrast, with reduced gradient strength, we found that the biasness of cellular migration goes down, as shown in Fig.3c (Video S7). Correspondingly, the average TI also significantly reduced to 0.28 ± 0.08 from the earlier TI of 0.44 ± 0.06 (Fig.2d).

To confirm that the observed migration is not caused by a rigidity gradient, we measured the elastic modulus (E) of the substrate along the gradient using atomic force microscopy (AFM). We found no gradient in elastic modulus (Fig.S4). We confirmed the uniformity of collagen density on the gel by immunostaining, thus eliminating the possibility of haptotaxis (Fig.S5). Further, to check if viscotaxis is ECM specific, we analyzed cell migration on another ECM i.e. laminin coated gel and found no significant difference (Fig.S6) (Video S8).

### Viscotaxis is driven by acto-myosin contractility

To understand the mechanism of viscotaxis, we first measured cellular traction on uniform Low Loss and High Loss substrates using Traction force microscopy (TFM). We found that the cells on Low Loss substrate apply ∼5 times more force than they do on High Loss substrate (Fig.4 a-c). This difference in traction may result in a force asymmetry on a substrate with loss modulus gradient resulting in the biased displacement of cells causing observed viscotaxis.

To verify this hypothesis, we inhibited actomyosin contractility by using two specific pharmacological inhibitors, blebbistatin and Y27632. Blebbistatin is known to inhibit myosin II^25^ and Y27362 reduces the cellular contractility by inhibiting ROCK^26^.

After 4 h of seeding on the substrate, hMSCs cells were treated with 20 μM blebbistatin and 10 μM Y27362. In the presence of the inhibitor, time lapse microscopy was performed for 18 h with 30 min interval. Fig.4 d-f represent rose plots for cell migration on substrate with loss modulus gradient in the presence of blebbistatin or Y27632. For control, only growth media and no inhibitor was used. We observed that the biasness in migration significantly drops in the presence of traction inhibitors. The same is reflected in the tactic index (TI) as shown in Fig.4g. In the presence of contractility inhibitors, the TI came down to value 0.07 ± 0.03 for blebbistatin and 0.09 ± 0.08 for Y27362 signifying no preferred direction for migration (Video S9, S10). As the cellular motility itself is dependent on contractility, we ensured that the concentration of the inhibitors used in this study reduced the cellular traction (data not shown) but did not affect cell velocity (Fig.4h).

**Fig.4.**
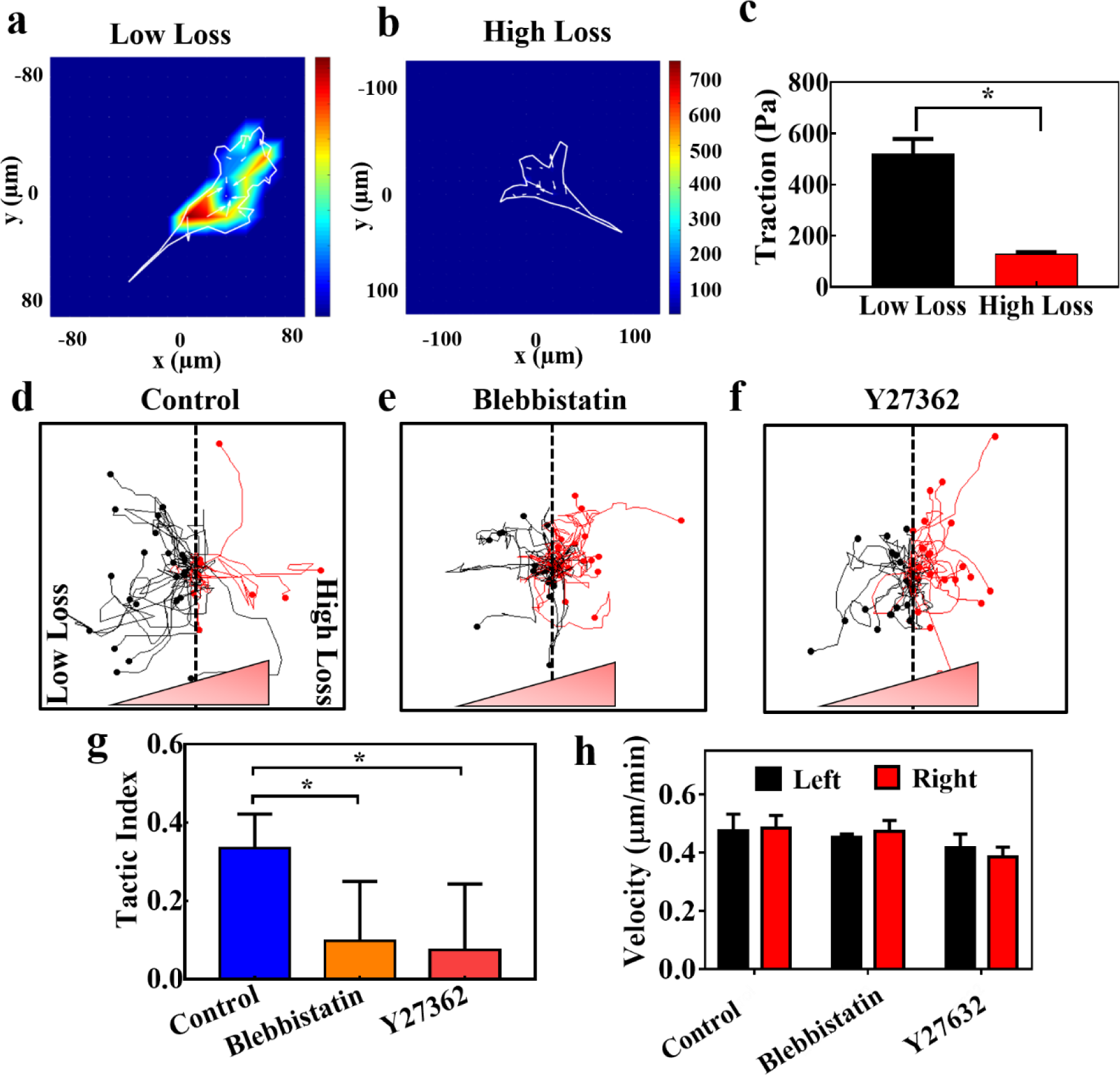
Viscotaxis is mediated through actomyosin contractility: Traction force applied by hMSCs on (a) Low Loss substrate, and on (b) High Loss substrate, (c) hMSCs on High Loss substrate showed lower traction than that of Low Loss substrate (n = 10 cells, * p< 0.05). Representative rose plots depicting trajectories of migrating cells for (d) without inhibitor (n = 106 cells). (e) blebbistatin (20 µM, n = 56 cells), and (f) Y-27632 (10 µM, n = 62 cells). (g) Tactic index without inhibitor, with blebbistatin, and with Y-27632. (h) Cell velocity after treating with blebbistatin and Y27632 compared with a cell with no drug treatment.

### Viscous substrate influences the cell membrane boundary of less motile cells

As a viscoelastic material shows creep behaviour, we predicted that a cell on such a substrate would not be able to apply traction in a sustained manner. If that is true, the fluctuation in traction should manifest in the cell spread area as well. However, testing this hypothesis with hMSCs is difficult as hMSCs are motile on both substrates. Thus, for this experiment we selected less motile HeLa cells and monitored the membrane boundary over 4 h with 5 min intervals. We observed that on High Loss substrate, cell boundary fluctuates more as opposed to the cells on Low Loss substrate where the boundary remains stable (Fig.5a,b)(Video S11,S12). In order to quantify this observation, cell boundaries were identified using image segmentation techniques in MATLAB. Furthermore, to avoid any effect due to cellular migration, these boundaries were overlapped to keep the centroid of the cell stationary (Fig.5c,d). In each image, the linear distance of the membrane from its centroid was measured for the 8 angular positions covering 360°. To reduce the noise in these measurements, distances along a ± 10° arc around each angle were averaged. These averaged radial distances were plotted with time for each cell (Fig.5 e,f). Standard deviations in these distances were calculated for each angle, and the 8 deviations were averaged to obtain an overall standard deviation, corresponding to the membrane fluctuations in one cell. This parameter was tested for statistical significance between the cells on Low Loss and High Loss substrate. We found that the overall deviation was significantly higher for cells plated on High Loss compared to the cells on Low Loss (Fig.5g).

**Fig. 5.**
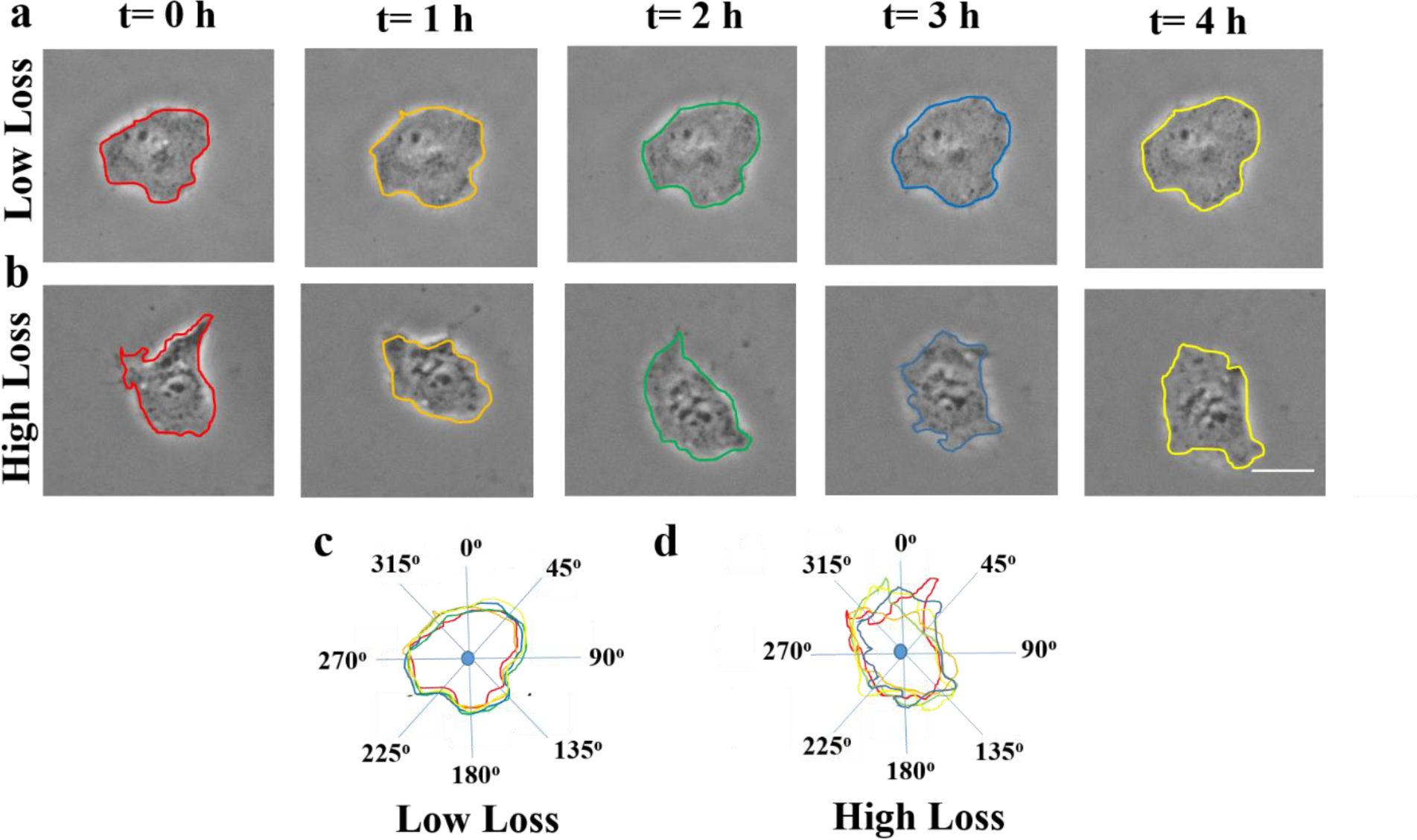
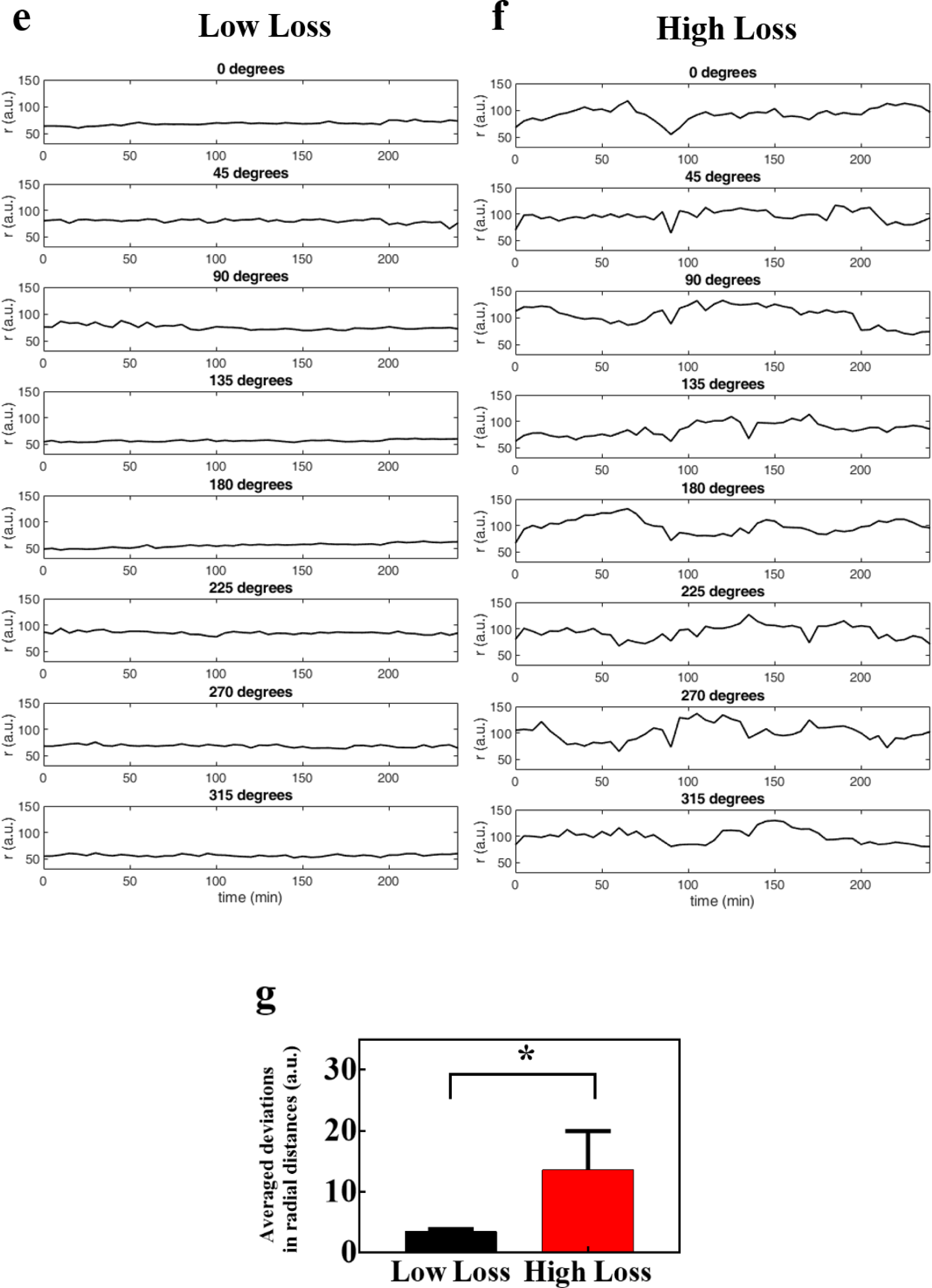
High loss substrate influences the cell membrane boundary of less motile cells: Time lapse of the boundary detection of a cell cultured on (a) Low Loss and, (b) High Loss substrate. (c, d) Merged boundaries of cells on Low Loss and High Loss with lines drawn with a reference point (nucleus) intersecting the membrane boundaries. Radial distance (r) with time plotted for 8 angles for one cell cultured on (e) Low Loss substrate and (f) High Loss substrate. (g) Student’s t-test showed a statistically significant difference between averaged deviations of radial distances in Low Loss and High Loss substrate, *p < 0.05 scale bar 100 μm; a.u.- arbitrary unit.

## Discussion

Tissue microenvironment *in vivo* is not purely elastic but viscoelastic in nature. Whether it is a cell, or ECM, or the whole organ, all relevant components of an organism have their own characteristic storage moduli (G’) and loss moduli (G”) ^27–29^. However, our understanding of the effect of viscoelasticity on different cellular behaviours including migration is nascent and incomplete. In this work, we have demonstrated that hMSCs migrate from high loss modulus region to low loss modulus region when exposed to a substrate with a gradient of G” but constant G’. We termed this so far unreported migration as “viscotaxis”. Viscotaxis is not a new term. This word was previously used to describe the preferential swimming behavior of non-mammalian cells in a liquid medium with viscosity gradient^31^. However, this is the first work to report the effect of solid viscoelasticity on the crawling migration of adherent cells.

There are two other cellular migrations or “taxis” well reported in literature namely durotaxis and haptotaxis that depend on adherent cues such as the gradient of rigidity and ligand density respectively^12, 32^. In this report, we created the substrates by varying the ratio and concentration of acrylamide and bis-acrylamide which may potentially cause a change in substrate rigidity and ligand presentation. To substantiate that the observed migration in this paper was not durotaxis or haptotaxis, we confirmed the uniformity of rigidity and the concentration of the ligand by AFM and immunostaining respectively.

Our result opens up an important question in the context of durotaxis. In all previous studies exploring durotaxis, researchers only considered the elastic modulus (E or G’). The changes in E or G’ were achieved by changing the polymer concentration or cross-linking density^33–35^. However, the same process may also change the substrate viscoelasticity and may have a pronounced effect on observed cellular migration. Our results suggest that any study probing the effect of substrate material properties on cellular migration must take the possible effect of substrate viscoelasticity into account. Our data also opens up the possibility to reexamine the published data on durotaxis and evaluate if the observed migration was a result of both the cues. Further study is needed to assess the combined effect of substrate storage and loss modulus on cellular migration.

We hypothesize that viscotaxis originates from the differential response of elastic and viscoelastic material to the same applied force. While an ideal elastic material attains a constant steady deformation instantaneously, a viscoelastic response is a time dependent phenomenon. If the same amount of stress is applied for the longer duration, the deformation increases with time; this phenomenon is known as creep. We explained viscotaxis with the help of creep and proportionality of cellular traction to the substrate resistance, as shown in Fig.7. When an adherent cell applies contractile force on a gradient substrate, the material respond in proportion to the applied stress. However, while at the more elastic end of the cell, deformation reaches a stable constant value, the other end continues to deform or creep (dashpot deformation, t= t0, t= t1 Fig. 7). As the resistance from the material decreases, cellular traction also goes down at that end causing a symmetry breaking, an essential condition for directed cell migration. Our hypothesis is based on three observations. One, cellular traction on the viscoelastic uniform gel is significantly lower than the same on the uniform elastic gel. This observation supports our hypothesis of traction asymmetry in a cell on a gradient gel. Two, if our predicted model is correct then a less migratory cell should show more dynamic behaviour in its spreading when seeded on a viscoelastic substrate than on an elastic one. We confirmed this with HeLa cells (Fig.6). Three, according to this model, viscotaxis is dependent on actomyosin contractility. We indeed found that viscotaxis gets impaired when treated with either blebbistatin or ROCK inhibitor Y27362. Both of these molecules are known to reduce cellular traction. A point to note here is that we used a minimal concentration of inhibitors that do not interfere with cell velocity.

**Fig. 7.**
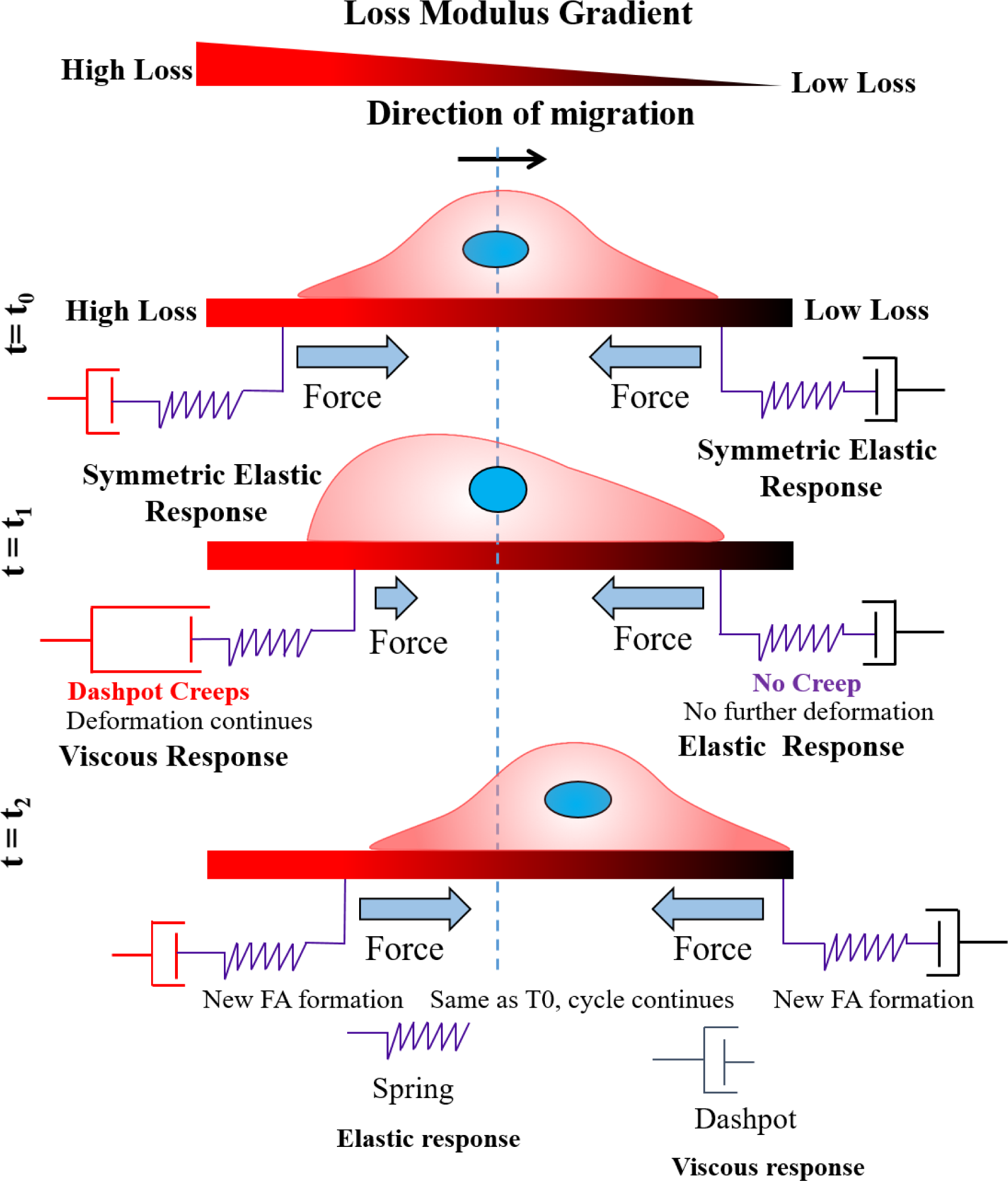
Schematic for loss modulus biased cell migration. Viscotaxis: cell migrates from High Loss to low loss modulus region. Gradient substrate with constant storage modulus (elastic) but varying loss modulus (viscous) component.

The bias during migration, as defined by the tactic index (TI), increases with an increase in gradient strength but was unaffected by the absolute value of G” for the range we tested (Fig.S4). Though both durotaxis and viscotaxis depend on relative material deformation and resulting asymmetry, durotaxis was reported to depend on both the gradient strength and absolute value of local rigidity. To answer whether viscotaxis is indeed independent of absolute loss modulus, one needs to check a wider range of G”.

## Conclusion

In conclusion, we have demonstrated that the gradient in loss modulus can be an important mechanical cue governing cell migration. We propose that viscoelasticity of the materials should be taken into account in the discussion of mechanotaxis. Further, its role in physiological and pathological conditions should be investigated in more details.

## Material and Methods

### Substrate preparation

Gels were formulated from base solutions of 40 wt% Acryl (Sigma-Aldrich) and 2 wt% Bis (Sigma-Aldrich). After mixing the solutions at different ratios in milli-Q water, Low Loss gel were fabricated by crosslinking 1 mL of solution with 1.0 μL of tetramethylethylenediamine (TEMED) (Sigma-Aldrich) and 10 µL of 10 wt% ammonia persulphate (APS) (Sigma-Aldrich). High Loss were fabricated by crosslinking 1 mL of solution with 1.5 μL of TEMED and 5 µL of 10 wt% APS. The gels were allowed to crosslink for approximately 15 min. The gradient gels were prepared using two drop diffusion technique as described earlier^12, 25^. Briefly, we made gradient gels by placing two drops (100 µl each) of High Loss and Low Loss solutions on a 22 mm * 22 mm square coverslip coated with di-chloro-dimethylsilane (DCDMS) of (Sigma-Aldrich). FluoSpheres (Invitrogen) 0.2 µm in diameter were added into the High Loss solution of 0.1% by weight in the gel solution. For shallow gradient gel preparation, hydrophilic cover slip functionalized with 3-(Mercaptopropyl) trimethoxysilane (MPTS) (Sigma-Aldrich) was placed on top of the drops, allowing the solutions to diffuse. For step gradient generation, gel solutions were allowed to partially polymerize for 5 min before placing the hydrophilic functionalized coverslip. In both cases, gradient gels were allowed to polymerize for 15 min.

### Gel mechanical testing

The viscoelastic spectrum (storage modulus, G’ and loss modulus, G”), as a function of frequency, of each of the resulting gels were measured using a rheometer (MCR301, Anton Paar) Solutions containing varying compositions of Bis and Acryl monomers were cross-linked between the plates of the rheometer lower plate. Both plates were roughen with polish paper (3 MM safety walk) and a stainless steel flat plate of 50 mm diameter was used with a gap distances 1mm. A time sweep test was conducted during the reaction to confirm that cross-linking had come to completion. The duration of the time sweep was 25 min with a controlled absolute strain of 1% and at an angular frequency of 1 Hz. Next, a frequency test was performed on the gels where the angular frequency was increased from 0.01 rad/s to 100 rad/s at a strain of 1%

### Gel functionalisation for cell seeding

The substrates were crosslinked with 100 µl of 5 µg/ml N-sulfosuccinimidyl-6-(4azido-2- nitrophenyl amino) hexanoate (Sulfo-SANPAH) (G-Bioscience) in HEPES buffer (pH=8.5) using UV cross-linker (Genetix) (312 nm) for 20 min. PAA gel substrates were then washed with Dulbecco’s Phosphate Buffer Solution (DPBS) twice to remove free molecules of sulfo-SANPAH and submerged in the collagen (25 µg/ml) or laminin (50 µg/ml) solution and incubated overnight at 4°C. Finally, before cell seeding, the substrates were kept in laminar hood under UV for 20 min following the washing with DPBS.

### Cell Culture

Umbilical cord human Mesenchymal stem cells (UC-hMSCs) were cultured in Dulbecco’s Modified Eagle Medium (DMEM) (Himedia) supplemented with 16% fetal bovine serum (FBS) (Himdeia), 1X Antibacterial-Antimycotic (Himedia) solution and 1X Glutamax (Invitrogen). UC-hMSCs were a kind gift from Prof Jayesh Bellare, IIT Bombay. Cells were passaged at around 70% confluency using 0.5% Trypsin-EDTA (Invitrogen) for 3 min at 37°C to detach the cells and centrifuged at 1000 rpm for 5 min. All experiments are performed with UC-hMSCs between passages 4 and 7. UC-hMSCs were seeded on the gels at a seeding density of 1000 cells per cm^2^. The cells were allowed to adhere to the gel and spread for a period of 4-5 h before time lapse microscopy.

### Time Lapse Microscopy

The time-lapse microscopy was performed using the Evos FL-Auto microscope (Life Technologies) attached with controlled environment on-stage incubator. Time-lapse imaging was started after 4 h of cell seeding and continued for 18 h duration with 30 min time interval between two consecutive images. The images were taken in two modes, first ‘PHASE’ mode to observe the cells, second in RFP mode to observe red fluorescent beads that determine the gradient boundary (Fig.S3).

### Cell migration analysis

ImageJ software (National Institutes of Health, Bethesda, USA) plugin with manual tracking (Fabrice Cordelières, Institut Curie, Orsay, France) was used to track the path of the cells from the images obtained over an 18 h period. During analysis on average 50-60 cells were tracked per substrate. Exported ASCII from manual tracking was imported to the software tool and the cell trajectories were all extrapolated to (X,Y) = (0,0) at time 0 h. Cell migration images were taken by setting up multiple sequential beacons that cover the gel along the gradient length. To analyze the Tactic Index, we needed to distinguish the cells migrating towards left and cells migrating towards right. This distinction was performed using plugin “Chemotaxis and Migration Tool” (IBIDI Software)^36^.

### Atomic force microscopy (AFM)

In Bio-AFM (MF3D, Asylum research), the probe of cantilever consists of micron sized tip. Cantilever deflection is used to calculate the indentation force. When the sample is indented by the probe, cantilever deflection is measured as a function of the probes z position. Thermal fluctuation was done in order to get the actual spring constant of the probe. To calculate the elastic modulus of the substrate, the indentation force was fitted to the Hertz model.

### Inhibitor Studies

hMSC cells were cultured on Low Loss and High Loss substrates at a density of 10,000 cells/cm^2^. Cells were allowed to spread for 4 h and then inhibitors such as blebbistatin (20 μM, Sigma) and Y27632 (10 µM, Sigma) were added and it was maintained for the entire duration of the respective studies.

### Traction Force Microscopy (TFM)

Gels for Traction Force Microscopy were prepared in a two-step process. High Loss substrate and Low loss substrate were made on 22 mm*22 mm^2^ coverslips. Gels were allowed to solidify, then 25 µl of High Loss and Low Loss solution having 1µm fluorescent beads (Fluka with a final concentration of 1:50) along with APS and TEMED was added on the hydrophobic plate, and then solidified gel was inverted onto the top of it and allowed to solidify. The gels were then treated with sulfo-SANPAH and coated with Collagen type-I as mentioned above. After 24 h of cell seeding, cells were lysed using Triton-X (1:100) without disturbing the gels, images of cells in a phase were taken before adding Triton-X, and the images of fluorescent beads were taken before and after Triton-X using EVOS FL Auto cell imaging system (Life Technologies). The code from J. P. Butler was used to calculate the bead displacement and traction force.

### Collagen staining

The 25 µg/ml collagen coated on uniform and gradient gels substrates were washed with DPBS for 3 times. These gels were incubated in custom made anti-collagen antibody/serum (gifted by Prof. Shamik Sen, IIT Bombay) (1:50) raised in rabbit for 4 h at room temperature. After incubation, gels were washed with DPBS twice, 10 min each and incubated with Alexa fluor 488 goat anti rabbit (1:500) secondary antibody for 3 h at room temperature.

### Cell boundary identification

Image segmentation techniques in MATLAB were used to obtain cell periphery from phase contrast microscopy images. Each grayscale image was cropped around the target cell, and the contrast was enhanced using contrast-limited adaptive histogram equalization (CLAHE)^37^ to increase edge-detection accuracy. Edge detection was performed using the Sobel algorithm, a gradient-based edge detection method^38^. The Boolean output consisting of image edges was then dilated by two pixels in each of the 4 cardinal directions and a hole-filling algorithm was applied to obtain a closed region over the cell. Next, image cleaning algorithms were implemented to remove all objects not pertaining to the target cell. Finally, the region was eroded by two pixels to compensate for the dilation step performed earlier, thus giving us the segmented region consisting of the cell. The periphery of this region overlapped well with the cell membrane and was used for statistical analysis. The centroid of the cell was obtained by averaging the x and y coordinates of the segmented region.

### Statistical analysis

Data are presented as means ± standard error of the mean and were analyzed using the Microsoft Excel software. Unpaired student t-test was for two samples and the one-way ANOVA for three samples. All test were performed assuming the non-gaussian distribution of the sample values. Statistical significant differences were defined as: * = p < 0.05 otherwise stated.

## Supplementary Materials

Figures S1-S6

Table S1

Videos S1-S12

## Acknowledgement

This research was funded by the Wellcome Trust-DBT India Alliance (Project # IA/E/11/1/500419), seed grant from IITB (Grant # 14IRCCSG002). We thank MHRD, IIT Bombay for providing the fellowship to PS and the Bio-AFM facility. We thank Dr. James P Butler (Harvard Medical School, Department of Medicine, Boston) for his TFM codes used for the analysis. We thank Shital Yadav for her valueable inputs.

