## SupplementaryMaterials for "“Viscotaxis”- Directed Migration of Mesenchymal Stem Cells in Response to Loss Modulus Gradient"

### Supplementary Information

| Sr. No. | % Acryl | % Bis | Storage modulus (kPa) | Loss Modulus (Pa) |
|---|---|---|---|---|
| 1 | 5 | 0.01 | 1.2 | 10 |
| 2 | 6.68 | 0.06 | 1.2 | 30 |
| 3 | 8 | 0.06 | 3.0 | 10 |
| 4 | 8 | 0.26 | 1.5 | 289 |
| 5 | 11.4 | 0.1 | 2.6 | 100 |
| 6 | 13.2 | 0.3 | 3.5 | 236 |
| 7 | 21.4 | 0.01 | 2.71 | 250 |
| 8 | 21.4 | 0.03 | 4.5 | 275 |
| 9 | 21.4 | 0.05 | 5.3 | 357 |
| 10 | 21.4 | 0.06 | 8 | 550 |
| 11 | 40 | 0.06 | 16 | 1400 |

**Table S1:** Polyacrylamide gel compositions and the resulting storage modulus (G') and Loss modulus (G") measured at 0.01 Hz and 1% strain.

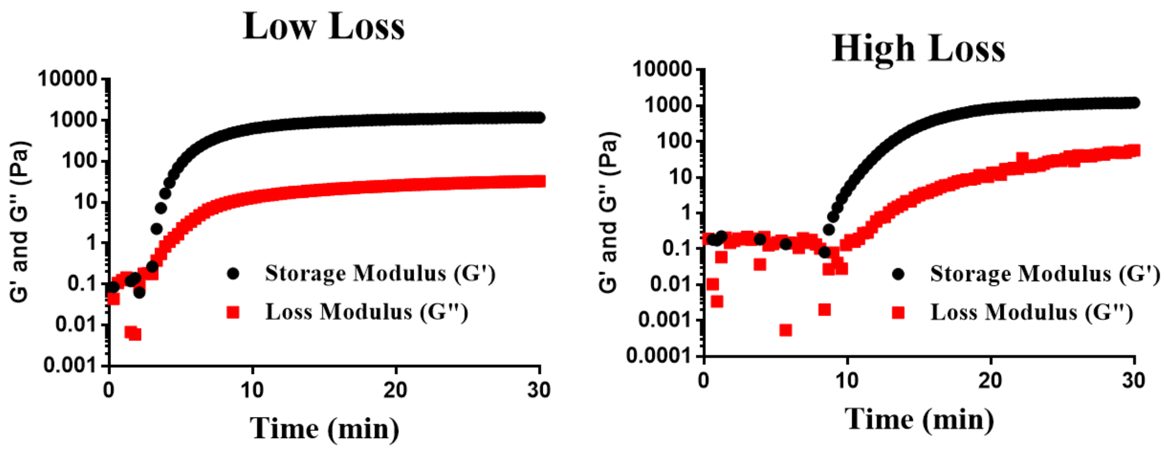


**Fig. S1.** Crosslinking and mechanical testing of polyacrylamide gels. Time sweep tests showing the development and plateau of Low Loss and High Loss subtrate storage moduli and loss moduli during crosslinking.

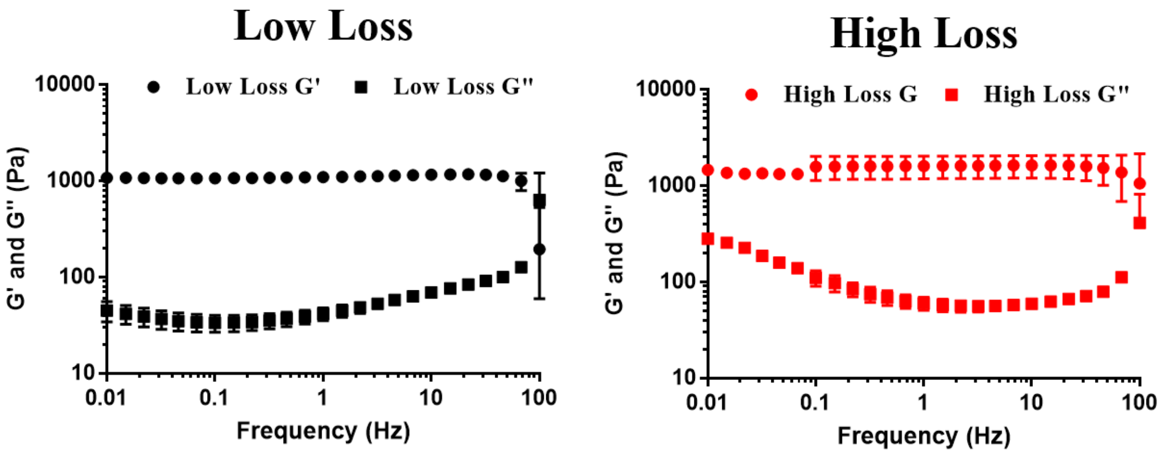


**Fig. S2.** Frequency sweep of Low Loss and High Loss gel samples performed with frequency decreasing from 100 to 0.01 Hz at a strain of 1 %.

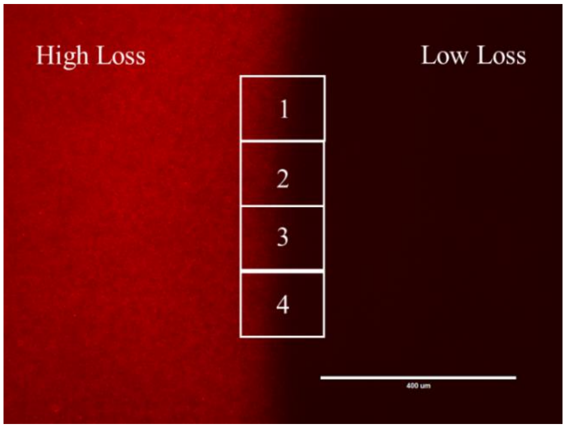


**Fig. S3.** The boundary of the High loss and Low Loss region of the loss modulus gradient gel: square boxes 1,2,3 and 4 at the boundary are the schematic representation of the the location choosen during the time lapse microscopy setting.

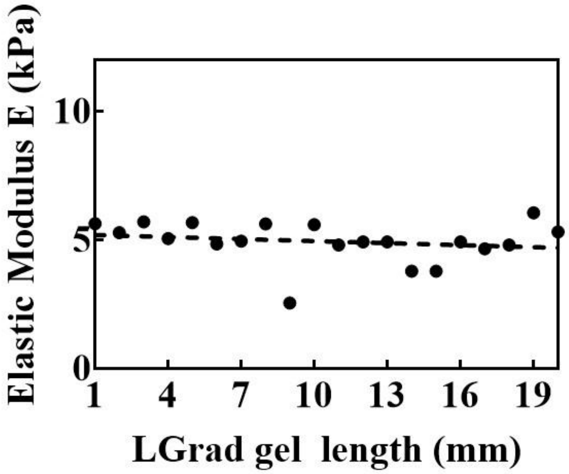


**Fig. S4.** Biased cell migration on loss modulus gradient is independent of elastic modulus. Elastic modulus measured across the shallow loss modulus gradient gels length (19 mm) using atomic force microscopy ( AFM ).

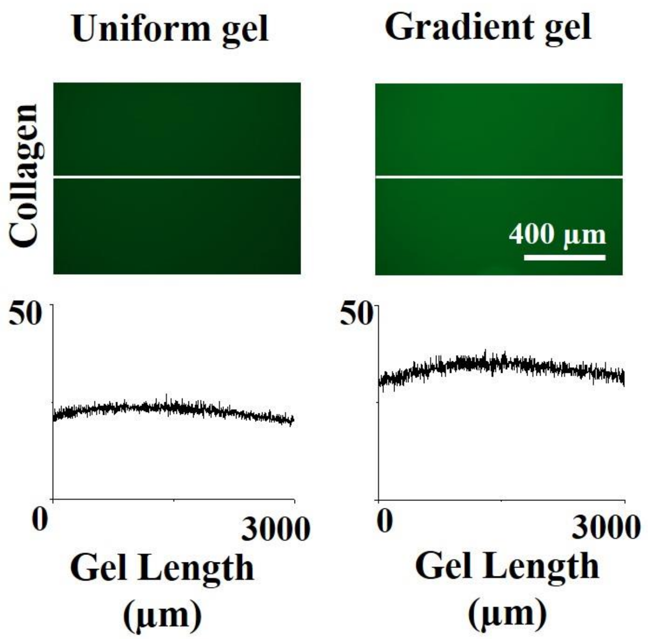


**Fig. S5.** Biased cell migration on loss modulus gradient is independent of collagen coating density. GFP Images of uniform and gradient substrate along with its corresponding image of GFP with anti-chicken IgG bound to primary anti-collagen antibody on collagen coated substrate.

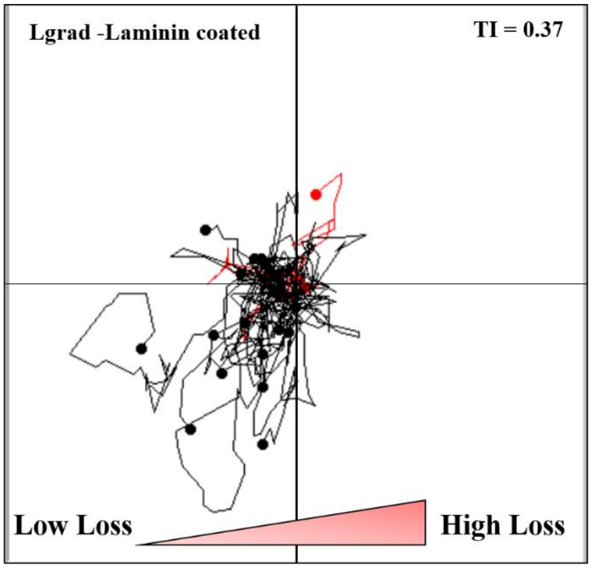


**Fig. S6.** Biased cell migration on loss modulus gradient substrate coated with ECM- Laminin. Representative rose-plots depicting the trajectories of migrating cells on 50 µg /ml laminin coated gradient of Low Loss – High Loss (shallow loss modulus gradient).

**Supplementary Videos**

**Video S1:** hMSCs migration on uniform low loss substrate.

**Video S2:** hMSCs migration on uniform high loss substrate.

**Video S3:** hMSCs migration on shallow loss modulus gradient coated with collagen

**Video S4:** hMSCs migration on loss modulus steep gradient. hMSCs migrating from high loss to low loss (event1).

**Video S5:** hMSCs migration on loss modulus steep gradient. hMSCs returning from high loss to high loss (event2).

**Video S6:** hMSCs migration on loss modulus steep gradient. hMSCs migrating from low loss to high loss (event3).

**Video S7:** hMSCs migration on loss modulus shallow gradient. hMSCs migrating from low loss to high (50:50 composition) loss.

**Video S8:** hMSCs migration on loss modulus shallow gradient coated with Laminin ECM. hMSCs migration from high to low loss modulus.

**Video S9:** hMSCs migration on loss modulus shallow gradient in presence of myosin inhibitor, blebbistatin (20 μm).

**Video S10:** hMSCs migration on loss modulus shallow gradient in presence of ROCK inhibitor, Y27632 (10 μm).

**Video S11:** Membrane fluctuation of HeLa cell on low loss modulus substrate

**Video S12:** Membrane fluctuation of HeLa cell on high loss modulus substrate.
